# Connecting tumor genomics with therapeutics through multi-dimensional network modules

**DOI:** 10.1101/083410

**Authors:** James T. Webber, Max V. Ranall, Swati Kaushik, Sourav Bandyopadhyay

## Abstract

Recent efforts have catalogued genomic, transcriptomic, epigenetic and proteomic changes in tumors, but connecting these data with effective therapeutics remains a challenge. In contrast, cancer cell lines can model therapeutic responses but only partially reflect tumor biology. Bridging this gap requires new methods of data integration to identify a common set of pathways and molecular events. Using MAGNETIC, a new method to integrate molecular profiling data using functional networks, we identify 219 gene modules in TCGA breast cancers that capture recurrent alterations, reveal new roles for H3K27 tri-methylation and accurately quantitate various cell types within the tumor microenvironment. We show that a significant portion of gene expression and methylation in tumors is poorly reproduced in cell lines due to differences in biology and microenvironment and MAGNETIC identifies therapeutic biomarkers that are robust to these differences. This work addresses a fundamental challenge in pharmacogenomics that can only be overcome by the joint analysis of patient and cell line data.

## INTRODUCTION

Cancer is caused by molecular aberrations that lead to de-regulation of cellular networks. Large-scale tumor sequencing efforts have sought to identify these molecular events in common cancer types across thousands of patients^1^. For example, The Cancer Genome Atlas (TCGA) cataloged the molecular aberrations present in human tumors by systematically measuring somatic mutations, copy-number alterations, gene methylation, transcriptomes and proteomes in over 11,000 tumors^2,3^. While these studies have been highly successful, the precise function of most of these molecular events is unclear, and exploiting them to better tailor therapies for patients remains a challenge. Computational approaches can help by annotating tumors based on pathway-level aberrations^4–6^, connecting tumors with relevant cell line models^7,8^ and grouping tumors that have similar global molecular phenotypes^2,9–13^. Such approaches have shown the importance of using the complementary nature of different profiling platforms to identify active cancer pathways. Thus far, such integration has been limited to curated pathway knowledge rather than the discovery of new functional networks^4–6^. The identification of pathways using a more unbiased approach can reveal new functionally related gene sets in cancer, which we refer to as gene modules.

In breast cancer, the discovery of molecular biomarkers has centered on classification and treatment based on gene expression subtypes^9,13–16^. The major subtypes in breast cancer include luminal A/B that display characteristics of cells of the breast lumen and are usually estrogen receptor positive (ER+), basal which share similarity to cells of the basement membrane and usually receptor negative and those that express the HER2 receptor^9^. However, patients often do not respond to therapies thought to be specific for their subtype^17,18^ and studies of drug sensitivities in breast cancer cell lines have shown that responses to 69-75% of drugs cannot be predicted by subtype^14,19^. This suggests that therapeutic efficacy is often based on the presence of molecular modifiers that are not represented by the global measures of subtype, cell type and receptor diversity.

To uncover biomarkers of drug response, several groups have profiled large collections of cell lines by measuring baseline molecular features as well as sensitivity to a wide range of compounds and using machine learning methods to identify therapeutic predictors^19–24^. While such approaches can generate statistically significant biomarkers in cancer cell lines, there have been challenges when translating them to human tumors^25,26^. One major challenge stems from the altered biology and tumor environment that distinguishes human cancers from cell lines^25,27^. Computational approaches that address this factor in the biomarker discovery process have not yet been developed.

In this paper, we present a new method called Modular Analysis of Genomic NETworks In Cancer (MAGNETIC) that integrates data across molecular profiling platforms by performing functional network analysis to identify tumor biomarkers and connect them to therapies. Using profiling data from breast cancers in The Cancer Genome Atlas (TCGA), we identify 219 gene modules that capture molecular features that are closely linked across samples and enriched for protein pathways. Using MAGNETIC we uncover new biological processes and present a quantitative landscape of micro-environmental factors in breast cancers that can inform new therapeutic targets. The method pinpoints molecular features that are limited to growth *in-vivo* and reveals that a surprising amount of gene expression and methylation data from human tumors reflects signals derived from the tumor environment that are not reflected in cell lines. Building on this finding, we show that modules preserved in cell lines can act as accurate biomarkers that are more robust than standard approaches because they are more reflective of a tumor context. This work reveals a new approach for the integrative analysis of molecular programs within human tumors and provides a powerful and clinically relevant way to connect tumor genotype to therapy.

## RESULTS

### Pathway signals are embedded across -omics platforms

We sought to measure the extent to which the interactions found in pathway databases carry information for interpreting correlations between molecular changes detected in cancer. We started with molecular profiles of primary breast cancers from the TCGA obtained using gene expression, DNA methylation, copy-number alteration, exome sequencing and Reverse Phase Protein Array (RPPA) platforms covering 941 patients^3^. After normalization, we constructed a correlation network by comparing all pairs of gene features across patients both within and between platforms. To measure concordance with known pathways we compared the distribution of these correlations to a compendium of 60,194 functional protein-protein interactions (PPIs) from the HumanNet database^28^.

We observed strong agreement of molecular signals between protein pairs known to interact both within and across –omics platforms in the TCGA. For example, two related ubiquitin specific peptidases USP32 and USP6 were not only highly co-expressed (r=0.95, p=5.5×10^−268^) but the copy-number of USP32 was also strongly predictive of expression of USP6 (r=0.73, p=6.4×10^−84^) (Fig. 1a,b). Beyond this gene pair, other pairs of molecular features at this level of correlation were enriched by 9-fold to interact and the most highly correlated gene pairs were 100-fold more likely to interact than at random (Fig. 1c). In another example, the kinase LCK was not only highly co-expressed with its substrate LAT (r=0.88, p=1.4×10^−181^), but it was also co-methylated (r=0.58, p=3.3×10^−87^) and the expression of LCK was highly anti-correlated with the methylation of LAT (r=−0.68, p=1.6×10^−74^) (Fig. 1d,e). Gene pairs with this level of anti-correlation were 21-fold more likely to interact and the most anti-correlated pairs were over 100-fold more likely to interact (Fig. 1f). Other comparisons within and across platforms revealed a similar trend. For example, gene pairs that were highly co-expressed (r>0.7) were 40-fold more likely to interact (Supplementary Fig. 1). These trends were also insensitive to the choice of pathway database, as analysis using the iRefWeb database revealed similar patterns (Supplementary Fig. 2)^29^. Thus, co-variation networks encode pathway information and could be used to identify new functional relationships in cancer.

**Figure 1:**
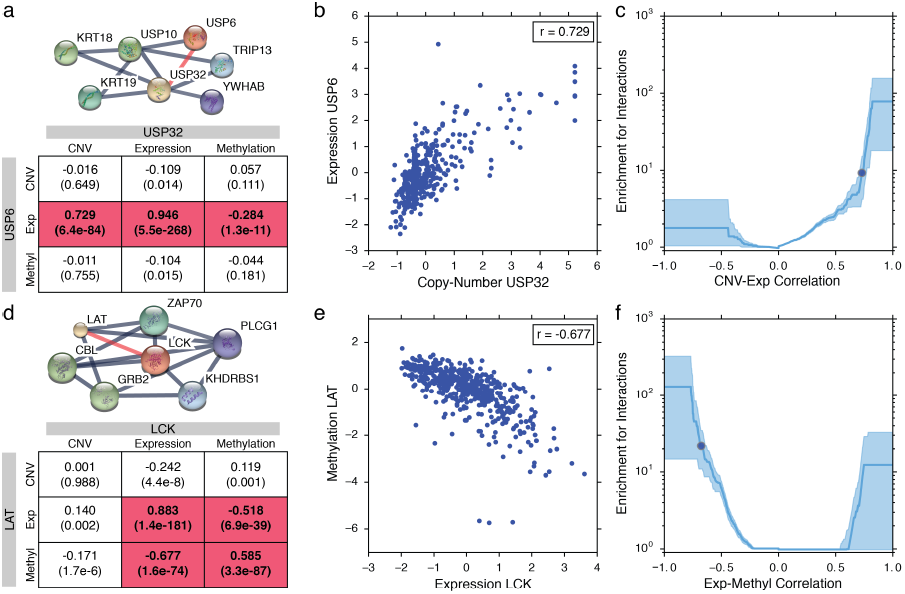
Pathway relationships are embedded in integrated molecular profiling data. **(a)** Interaction network of ubiquitin specific peptidases USP6 and USP32 from the STRING database and Pearson correlation between molecular features of USP6 and USP32 across TCGA breast cancers. P-value of association in parentheses and those that are significant after correction for multiple testing are highlighted (p≤0.05). **(b)** Scatter of normalized USP32 copy-number and USP6 expression across the TCGA. **(c)** Enrichment of protein interactions among gene pairs at a given correlation cutoff. Enrichment is calculated by comparison of the fraction of gene pairs at a given correlation cutoff that interact in HumanNet versus a random background (see Methods). For gene pairs that correlate ≥0.729, there is a 9-fold enrichment for known interactions (dot). Shaded areas reflect the 95% confidence interval. **(d)** The interaction network of the kinase LCK and its substrate LAT and relationships between their molecular profiles across platforms. **(e)** Scatter of LCK expression and LAT methylation. **(f)** Enrichment over random of gene expression and methylation correlations for gene pairs that interact in HumanNet. For gene pairs with a correlation of ≤−0.677 there is a 21-fold enrichment.

### A Modular Analysis of Genomic NETworks in Cancer (MAGNETIC)

Based on the observation that molecular correlations are reflective of pathway co-membership we sought to integrate these gene linkages into a single network. The multi-platform correlation network can be visualized as an undirected graph with multiple layers, each representing a data type (Fig. 2). Each gene corresponds to a node in every layer and edges connect nodes both within and between layers. There are 15 different edge types (all pairs of five data types, with repetition). Next, we re-scored the network edges by mapping a given correlation value to the log-likelihood ratio (LLR) of it reflecting a known interaction and merged them into a single network (see Supplementary Methods). The integrated network was highly enriched for known PPIs and outperformed any single correlation network type (Supplementary Fig. 3).

**Figure 2:**
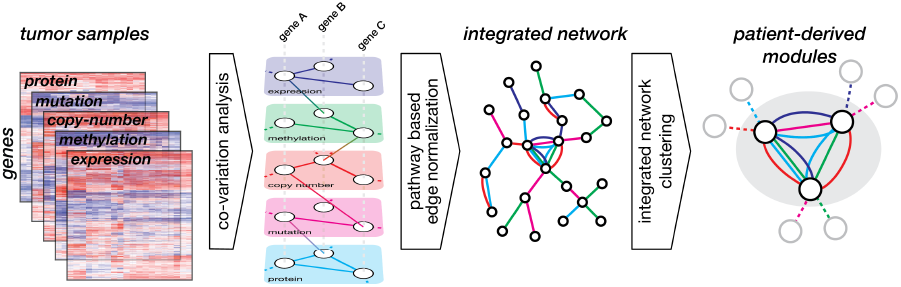
The MAGNETIC workflow. We begin with normalized DNA copy-number, methylation, somatic mutations, mRNA expression and protein abundance data from a collection of tumor samples. We compute a multi-layer gene similarity network by computing the correlation between all pairs of gene features both within and between profiling platforms. Each linkage in this correlation network is normalized through comparison against a benchmark of known protein pathways. Scored edges are then merged into an integrated network in which nodes represent genes and multiple edges between nodes represent co-incidence of different types of linkages. Clustering of this network using a random walk algorithm reveals gene modules whose components are closely related in multiple data types.

We next used a network-clustering algorithm based on repeated random walks to find densely interconnected modules within the integrated network^30^. Clustering of this network at different score cutoffs revealed an optimal overlap with known PPIs at an LLR>3 (Supplementary Fig. 4). At this cutoff, we identified a total of 219 modules with a median number of 18 genes per module and scored them in the TCGA cohort (Supplementary Fig. 5, Supplementary Table 1, 2). The modules were highly integrative with 84.5% reflecting data from multiple platforms. For example, the module containing ERBB2 (HER2) consisted of 25 genes and included 60 co-expression and 44 co-copy number variation edges at an LLR>3, reflecting the coordinated amplification and expression of genes in the HER2 amplicon (Fig. 3a,b). Module 37, based largely on gene expression, reflected the status of ER and was enriched for its direct transcriptional targets (p=3.1×10^−6^) (Fig 3c,d)^31^. Sixteen percent of modules were significantly enriched for a particular GO, KEGG, or Reactome pathway or function, and 44% contained known PPIs (Supplementary Table 1). Modules recapitulated previously published mutational events in breast cancers such as TP53 (#181), PIK3CA (#9), GATA3 (#37), and MAP3K1 (#5) as well as published gene signatures associated with proliferation, stromal involvement and angiogenesis^32,33^ (Supplementary Table 3). Therefore, we conclude that our approach integrates data from diverse platforms covering 48,093 unique gene and protein features into a set of 219 pathway-enriched modules that decompose complex tumor genomes into independent molecular signatures.

**Figure 3:**
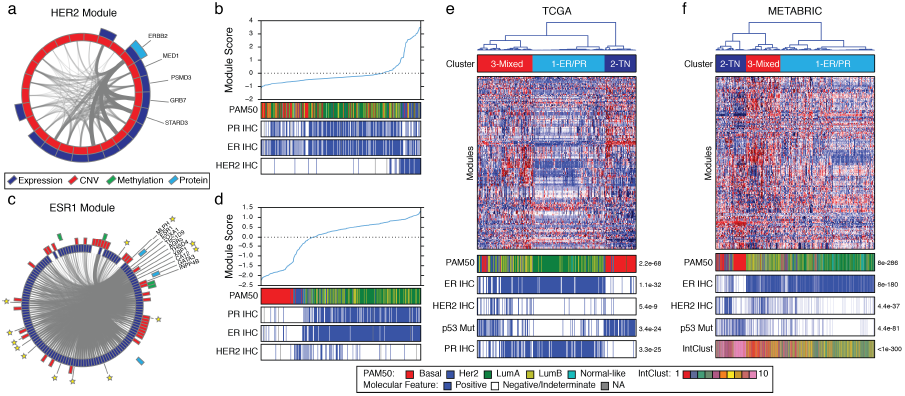
Gene modules recover known breast cancer biomarkers. **(a)** Circos plot representation of the module network containing HER2. Colors represent different data sources selected in the final integrated network for each gene and edge thickness is proportional to edge score. Top central genes are labeled. **(b)** TCGA samples sorted by HER2 module score. PAM50 subtype and molecular receptor status as determined by IHC are shown. **(c)** The module network containing the estrogen receptor, ESR1. Direct transcriptional targets of ER as assessed through ChIP analysis are marked with a star. **(d)** TCGA samples sorted by ESR1 module score. **(e)** Consensus clustering of 219 modules scored across TCGA breast cancers. Three subtypes and their association with subtype, IHC status for ER, PR, and HER2, and p53 mutational status are shown. **(f)** Consensus clustering of module scores in the METABRIC dataset identifies three clusters. Association between clusters and PAM50, IntClust subtypes, IHC markers and TP53 mutation are shown. P-values based on chi-squared test. TN=triple negative.

### An integrated module map of human breast cancers recapitulates known subtypes and markers

Because modules form a reduced representation of the molecular features in breast cancers, we next asked whether the set of module scores for individual tumors could be used to support published subtyping approaches in this disease^9,13^. Consensus clustering of all module scores across the TCGA cohort revealed the presence of three patient groups that largely supports the molecular subtypes called by PAM50 (p=2.2×10^−68^ via *χ*^2^ test), as well as the receptor status of ER (p=1.1×10^−32^), PR (p=3.3×10^−25^) and HER2 (p=5.4×10^−9^) (Fig. 3e, Supplementary Fig. 6a,b and Supplementary Table 4)^9^. Further interrogation revealed that Cluster 1 is largely ER/PR positive, Cluster 2 is largely triple-negative due to an absence of receptor expression and enrichment for TP53 mutations^3^, and Cluster 3 is mixed ER, PR and HER2 positive (Fig. 3e). As validation in an independent cohort, we scored modules based on available copy-number and gene expression data across 1,966 patients in the METABRIC study (Supplementary Table 2)^13^. Consensus clustering of this cohort also revealed three subtypes that mirrored those found in the TCGA (Fig. 3f, Supplementary Fig. 6c-e and Supplementary Table 4). In addition, our three subtypes coincided with receptor status and molecular subtypes identified in the original study using IntClust (Fig. 3f)^13^. This stratification based on receptor status largely parallels recent work showing that HER2+ tumors are not restricted to any particular subtype^34^. Therefore, we conclude that the modular decomposition of the TCGA allows for the grouping of patients with similar tumor phenotypes and similar disease progression characteristics largely in agreement with previous approaches.

### Modules reveal a role for differentiation via H3K27 tri-methylation in breast cancer

We observed a module (#27) that was largely based on expression which was highly enriched for genes whose promoters are marked by histone H3 Lysine 27 tri-methylation (H3K27me3) in ES cells, including SOX1, NEUROG1, NEUROG3, FOXB1 and FOXD3 (Fig. 4a,b). H3K27me3 is deposited by the Polycomb Repressive Complex 2 (PRC2) histone methyltransferase complex consisting of EZH1/2, SUZ12, RBBP7/4, and EED^35,36^. This modification is associated with the repression of genes involved in development and differentiation, but its role in breast cancer is almost completely unknown. To determine if genes in this module are regulated via H3K27me3 in breast cancer we obtained H3K27me3 ChIP-seq data for three breast cancer cell lines: SUM159PT (score=0.11), T47D (0.37) and MCF7 (1.39)^37^. We first verified that the module score reflected the expression level of genes within the module (Fig. 4c). We found that the promoters of gene in module 27 were more likely to be marked by H3K27me3 than other genes and that the level of H3K27me3 occupancy was higher in cells with a low module score indicating that histone modification state regulates the activity of this module (Fig. 4d, Supplementary Table 5). EZH2 over-expression has been implicated as an oncogene in many epithelial cancers^38^ and analysis of RNA-seq data from murine lung tumors driven by EZH2 over-expression indicated a specific down-regulation of genes in module 27 compared to non-transformed lung tissue (Fig. 4e)^39^. In support of a similar mechanism in breast cancer we found that EZH2 expression was anti-correlated with the activity of this module in breast cancer cell lines (r=−0.34, p=0.029) (Supplementary Fig. 7). Given the significance of PRC2-driven epigenetic regulation in other cancers, determining the importance of regulating this module during tumorigenesis could yield new biological insights and clinical targets for breast cancer.

**Figure 4:**
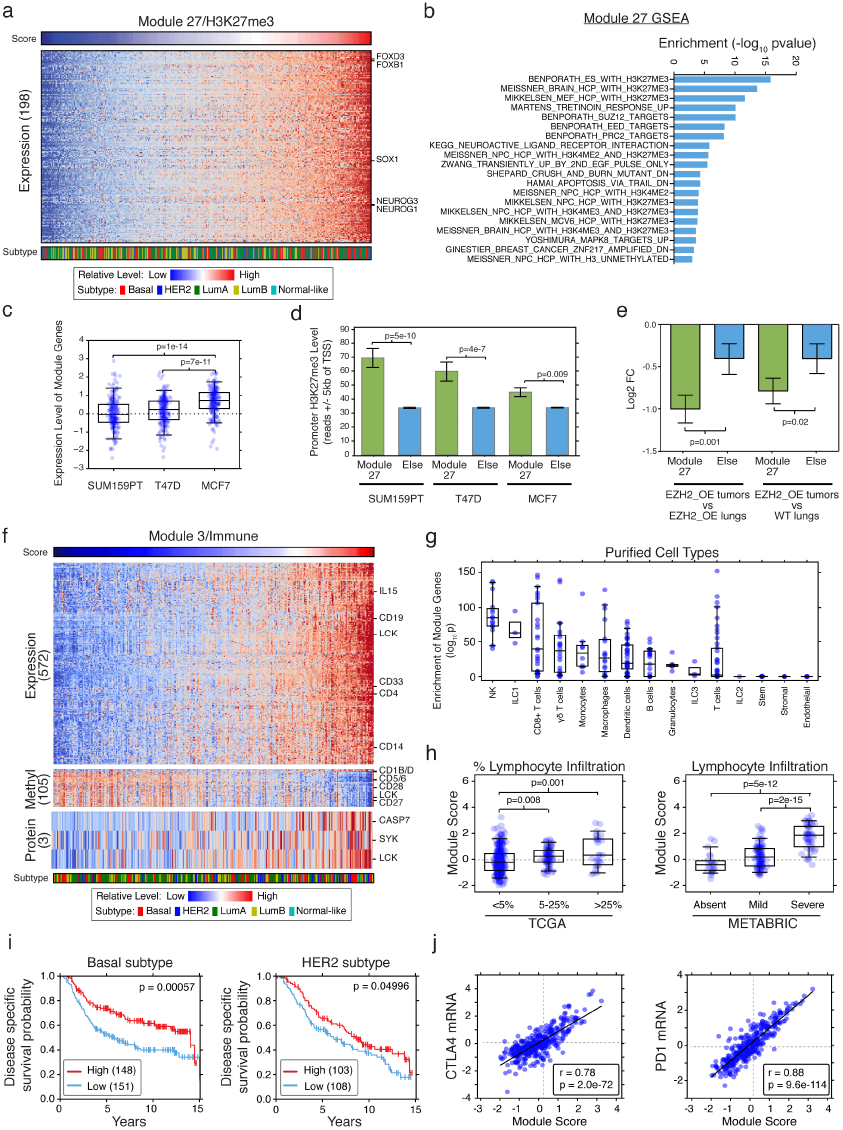
Modules that reflect coordination of gene expression by Histone 3 Lys27 tri-methylation and report on the tumor immune microenvironment. **(a)** Heatmap of molecular features associated with module 27 (r^2^>0.1). **(b)** Gene Set Enrichment Analysis of module genes. **(c)** Relative expression of module genes in SUM159PT, T47D, and MCF7 cell lines. **(d)** The number of reads in H3K27me3 ChIP-Seq samples within promoters of module genes compared to background in SUM159PT, T47D, and MCF7 cell lines. **(e)** Repression of module genes in EZH2 driven lung tumors as compared to EZH2 over-expressing lungs and wild-type lungs in an EZH2 genetic mouse model. **(f)** Heatmap of molecular features associated with the overall activity of the immune module (r^2^>0.1). For clarity, the CNV of one gene is not shown. **(g)** Enrichment for high expression of module genes from normalized RNA-seq data in 227 purified immune cell type datasets. Cell types are categorized into 15 groups and enrichment based on a t-test. **(h)** Comparison of module scores with annotated lymphocytic infiltration values in TCGA and METABRIC datasets. **(i)** Kaplan-Meier disease-specific survival curves of METABRIC samples stratified by median module score in Basal and HER2 tumors. Log-rank p-value shown. **(j)** Correlation of module score with expression of CTLA4 and PD1 in the TCGA.

### Modules as reporters of the tumor microenvironment

The tumor microenvironment exerts a high level of control on tumor behavior and can influence the interpretation of genomic analyses^40,41^. The influence of the microenvironment is apparent in the activity of several modules. Module #3 was highly enriched for genes related to the immune system (GO enrichment, p=1×10^−160^, Supplementary Table 1). This included the expression of B cell marker CD19, T cell marker CD4, and NK cell marker IL15, and protein abundance of the T cell kinase LCK (Fig. 4f)^42,43^. Comparing with gene expression data from purified cell populations we found that genes in the module were highly expressed in NK cells, ILC1 cells, T cells, monocytes and macrophages (Fig. 4g, Supplementary Table 6)^42^. Furthermore, we found that the activity of Module #3 was related to general lymphocyte infiltration scores based on pathological assessment in both the TCGA and METABRIC cohorts (Fig. 4h). Next, we sought to explore how this module might influence disease progression. This module was highest in the HER2 and Basal PAM50 subtypes (Supplementary Fig. 8) where it is associated with a favorable outcome (Fig. 4i), consistent with the positive role of T cell infiltration on cancer prognosis^44^. Interestingly, the module contained immune checkpoints PD1 and CTLA4, both markers of T cell exhaustion^45^ (Fig. 4j). Therefore, while infiltrating immune cells reflect anti-tumor recognition and promote a tumor suppressive environment, we hypothesize that these patients could benefit from re-activation of resident immune cells through anti-CTLA4 or anti-PD1 immunotherapy.

Other modules were reflective of non-tumor cell types. Module #12 enriched for a stromal gene signature and included many collagens associated with the extra-cellular matrix (ECM) (p=4.1×10^−52^, Supplementary Fig. 9a)^33^. This module was correlated with pathological assessments of stromal cells and indicated tumors with the presence of significant ECM involvement (Supplementary Fig. 9b,c). Module #16 was enriched for genes involved in vascularization and highly expressed in endothelial cells, including F10 and KDR/VEGFR2 (Supplementary Fig. 9d, Supplementary Table 6). Tumors scoring highly for module #16 were highly vascularized and less necrotic based on pathological assessment (Supplementary Fig. 9e,f). Taken together, these data indicate that gene modules report on the presence of components of the microenvironment that would be difficult to identify by approaches that are strictly limited to known pathways.

### Identification of preserved gene modules in cancer cell lines

We expected that modules reflecting tumor-specific variation would be more accurately modeled by cell lines in culture and therefore more useful in pharmacogenomics efforts. We therefore sought to exclude modules that reflect processes not captured in cell culture due to alterations in the growth environment. Since each module represents a set of relationships in an underlying molecular network, and we reasoned that modules which maintain these relationships in both tumors and cell lines are likely to reflect shared biology. We investigated whether the molecular features that are linked in the TCGA network were also linked in a panel of 82 molecularly characterized breast cancer cell lines^19^. For each module we calculated an edge preservation score reflecting the average increase in the pairwise correlation between molecular features when calculated across cell lines as compared to background (see Supplementary Methods). While there is no clear cutoff of edge preservation score, based on the resulting bimodal distribution of scores we chose a threshold of module preservation that excluded 59 modules to simplify further analysis (Supplementary Fig. 10). As expected, modules reflecting tumor specific events such as amplification of HER2 were identified as preserved (#92) and modules associated with the tumor microenvironment were not (#3 - immune, #12 - stromal, and #16 - endothelial) (Fig. 5a).

**Figure 5:**
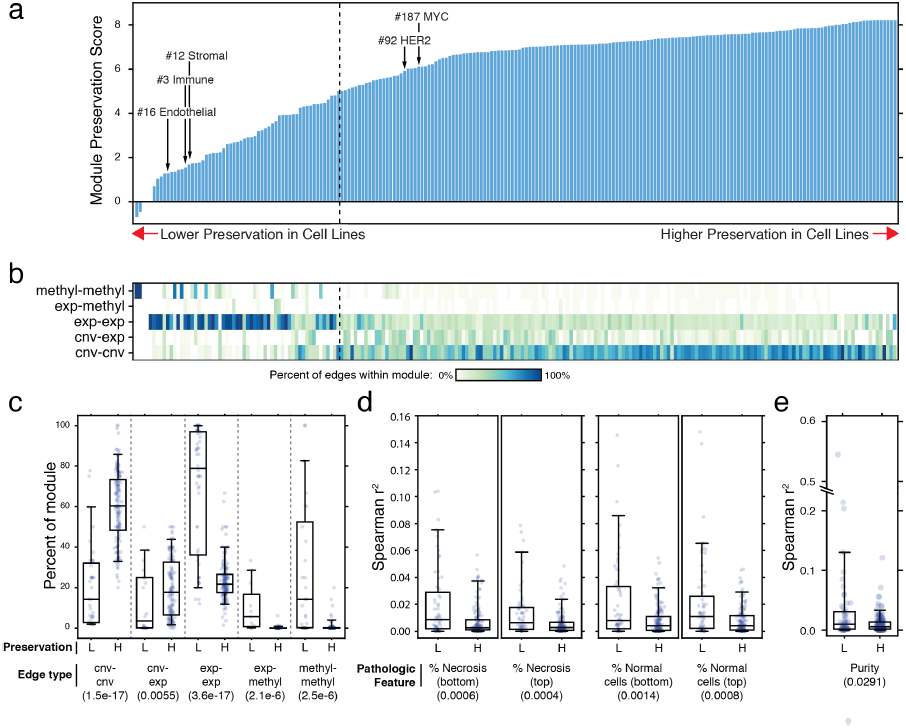
TCGA modules not preserved in breast cancer cell lines are associated with tumor impurity and necrosis. **(a)** Sorted preservation scores for 219 breast cancer modules evaluated in cell lines. Lower preserved modules have a score less than 5 (dotted line). **(b)** For each module in (a), the percent of the LLR>1 network that corresponds to each edge type are shown. **(c)** Percent of each edge type for lowly (L) and highly (H) preserved modules in the LLR>1 network. **(d)** Correlation of module scores with pathologic assessments of necrosis and normal cell infiltration for lowly and highly preserved modules. Top and bottom refer to different physical locations within the tumor. **(e)** Comparison of module types with computational assessment of tumor purity. P-values based on Mann-Whitney test in parenthesis.

The differences in module preservation led us to ask whether preserved modules were more likely to reflect data from certain types of –omics platforms. Modules associated largely with gene expression or methylation were much less likely to be preserved (Fig. 5b,c). In support, gene networks based solely on co-expression or co-methylation in the TCGA only marginally overlapped with such networks from breast cancer cell lines whereas networks that include copy-number variation (CNV) as a component were much more robust (Supplementary Fig. 11). There was no difference in the frequency of CNV-expression edges in highly versus lowly preserve modules, suggesting that expression data is most useful when tied to genomic events that are tumor specific. Overall, the activities of the 59 lowly-preserved modules were significantly associated with pathologic assessments of necrosis and normal-cell infiltration, as well as computational assessments of tumor impurity (Fig. 5d,e)^41^. Based on the 2,596 genes present in lowly-preserved modules we estimate that the expression and methylation of at least 13% of the genome reflects differences in biology between human tumor samples and cell lines, including the influence of the tumor microenvironment. Our analysis suggests that breast cancer biomarkers in cell lines tied to events such as copy-number variation and mutation are more likely to yield clinically translatable biomarkers because they are the most robust to issues such as tumor purity.

### A module-drug network identifies determinants of drug sensitivity that are robust to differences between patients and cell lines

We next investigated whether the preserved modules could be used for therapeutic stratification across a panel of 82 breast cancer cell lines profiled across 90 drugs^19^. We found a total of 271 module-drug relationships covering 74 drugs and 99 modules with an FDR<5%. We focused our attention on the 58 drugs (64.4%) whose response could not be predicted by subtype (Fig. 6a; Supplementary Table 7). While we identified known connections such as the HER2 module associated with sensitivity to the HER2 inhibitor Lapatinib (Fig. 6b), we also identified 97 connections among drugs where the module combined with subtype information was more predictive than subtype alone (Fig 6a,c).

**Figure 6:**
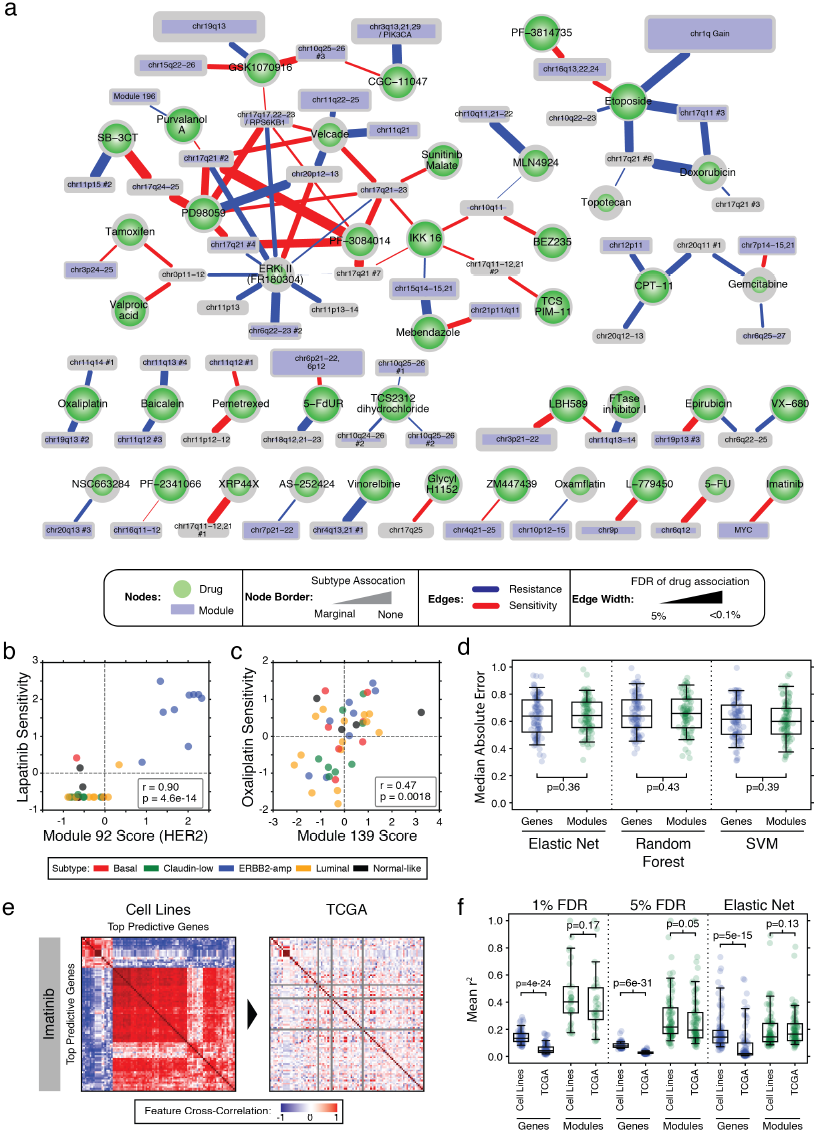
A Module-drug network identifies high performance biomarkers that are preserved between patients and cell lines. **(a)** Network of 97 module-drug associations based on breast cancer cell line modeling. Modules significantly associated with drug response are shown (FDR≤5%). Drugs are limited to those that are not associated with molecular subtype based on an FDR threshold of 5%. The size of each module is proportional to the number of genes within it. **(b)** Scatter plot of cell line association of Lapatinib response with module #92 (HER2) and **(c)** Oxaliplatin with module #139 (chr11q14#1). Cell lines colored by subtype. P-values based on Pearson correlation. **(d)** Comparison of median absolute error of cross-validated predictions of drug sensitivity using single gene features or modules as input to elastic net, random forest or SVM based predictors. P-values based on Mann-Whitney Test. **(e)** Cross-correlation for all pairs of molecular features that are the most predictive of response to Imatinib in cell lines at an FDR of 1% and cross-correlation of the same features in the TCGA. **(f)** The average cross-correlation (r^2^) of features selected by various statistical methods (FDR, elastic net) using single genes or modules in cell lines and evaluation of cross-correlation of the same features in the TCGA. Each point represents a model for a single drug. P-values based on Mann-Whitney Test.

Current pharmacogenomics approaches are based almost exclusively on data from cancer cell lines^19–21,23,24^. We reasoned that the highly preserved modules may constitute biomarkers that not only show strong performance as predictors of drug responses but also are more likely to translate when evaluated in human tumors. We first established that modules perform comparably to genes as features used to build predictive models of drug responses using common methods of machine learning for this task (Fig. 6d)^24^. Central to the performance of a predictive biomarker is the structure of the relationships between the features it uses. Therefore, to assess if a biomarker is likely to translate when applied to human tumors we examined the relationships among its features by measuring their cross-correlation independently in cell lines and in the TCGA. Surprisingly, we found that the relationships among features based purely on cell line data were completely altered in human tumors. As an example, top molecular features correlated with Imatinib sensitivity were highly cross-correlated in cancer cell lines (mean r^2^=0.165), but these relationships are completely lost in tumor samples (r^2^=0.015) (Fig. 6e). Logically, if the gene inter-relationships used to construct a biomarker are not maintained in tumors, it cannot be predictive of its intended drug response. We performed this analysis across all drugs using both gene sets and modules, using simple rank based methods (FDR cutoff of 1% and 5%) as well as with elastic net regression. Applied to all drugs, biomarkers based on genes had a significantly reduced cross-correlation in the TCGA when compared to cell lines, whereas module-based approaches maintained a consistently high cross-correlation in both cell lines and human tumors (Fig. 6f). We conclude that preserved modules reflect molecular relationships found in both cell lines and tumors and can aid in the development of improved, clinically relevant biomarkers.

## DISCUSSION

MAGNETIC integrates genomics, transcriptomic, epigenomic, and proteomic data across breast cancers to identify a set of gene modules that have coordinated activity across patients. As opposed to previous approaches for identifying gene modules based solely on correlation^46–48^, our approach integrates complementary data types based on comparison with known protein-protein interactions and the resulting modules show a strong enrichment for known pathways and shared functions. Many identified modules were concordant with disease subtypes, cell surface receptors and prognostic gene signatures. In contrast to previous approaches the method is not constrained to pathway definitions, which allowed the discovery of patterns in cancer data that are reflective of distinct cell types and epigenetic programs^4,49^. Future work could explore patterns of module activities across other disease types such as those found in a pan-cancer analysis of tumor genomes^2^.

Interrogation of the modules highlights an epigenetic program regulated by H3K27 tri-methylation and the presence of various micro-environmental cell types. The H3K27me3 module represents the activity of genes involved in differentiation that are repressed by the PRC2 complex. While the role of PRC2 activity in breast cancer maintenance is largely unknown, it is possible that tumors with low module activity would respond to inhibitors of the PRC2/EZH2 complex^39,50^. The identification of 603 genes reflecting immune cell infiltration also represents a potential avenue for therapeutic targeting. This finding is largely concordant with recent work showing that immune infiltration is positively linked with survival in ER negative breast cancer^13,44,51,52^. The co-incidence with markers of T-cell exhaustion suggests that this module may predict responsiveness to immune checkpoint blockade.

The development of biomarkers from cell line data is a subject of intense investigation in the field^14,19–21,23,53–55^. One powerful aspect of our approach is that it allows for the placement of genes into a putative molecular network (i.e. the module), and the preservation of this network in cell lines can be assessed.

We quantified the preservation of modules across a panel of breast cancer cell lines and found that many of the strongest co-expression and co-methylation signals in the TCGA were not preserved in cell lines. Our results indicate that the expression and methylation of at least 13% of the genome in TCGA samples reflect changes due to altered biology and the tumor microenvironment. Based on these issues, we propose that first learning a set of robust biomarkers from patient data and then evaluating them in cell lines could lead to increased success in biomarker development. We provide key evidence that extant procedures used to generate pharmacogenomic biomarkers are highly prone to error, as the relationships upon which they are built in cell lines fall apart when translated into human tumor specimens. As a solution, we develop a framework to use modules as the basis for biomarker discovery that shows comparable performance but is resistant to this source of error. We expect that computational data integration at two levels, the first between -omics platforms and the second between *in vivo* and *in vitro* samples, will ultimately aid in the translation of the cancer genome into clinical practice.

## ACKNOWLEDGEMENTS

The authors would like to thank Dvir Aran, David Quigley, Nevan Krogan, and Andrej Sali for helpful comments. This work was funded by U01 CA168370 and R01 GM107671 to S.B.

## AUTHOR CONTRIBUTIONS

J.T.W. and S.B wrote the manuscript and conceived the method. J.T.W. performed computational analysis. S.K. provided key datasets. M.V.R. provided essential insight and advice.

